# The Enhanced Activation of Innate Immunity in *Drosophila* S2 Cells by *Micrococcus luteus* is Mediated by Relish

**DOI:** 10.1101/2024.09.02.610512

**Authors:** Zaur M. Kachaev, Mona Ghassah, Anton A. Musabirov, Alexander V. Shaposhnikov, Ilya Y. Toropygin, Yulia A. Polunina, Nikita G. Stepanov, Victor K. Chmykhalo, Yulii V. Shidlovskii

## Abstract

The IMD and Toll signaling pathways in *Drosophila melanogaster* mediate the innate immune responses to Gram-negative and Gram-positive bacteria and fungi, respectively. Here we studied the involvement of the NF-κB transcription factor Relish, which is a mediator of the IMD pathway, in the humoral immune response to the Gram-positive bacteria *Micrococcus luteus* and *Bacillus subtilis* and the entomopathogenic fungus *Metarhizium anisopliae*, using *D. melanogaster* S2 cells as a model. Activation of Relish proteolysis was observed after S2 cell treatment with the control Gram-negative bacterium *Escherichia coli*. We found that *M. luteus* had also a noticeable effect on Relish activation, while *B. subtilis* and *M. anisopliae* effects were modest. Activation patterns of the genes encoding predominantly the IMD-pathway-dependent antimicrobial peptides (AMPs) and peptidoglycan recognition proteins (PGRPs), as well as the levels of Relish recruitment to the promoters of the genes, were found to be very similar in S2 cells treated with *E. coli* or *M. luteus* but were lower and differed in the case of *B. subtilis* and *M. anisopliae.* A Relish knockdown (KD) decreased the induction levels observed for all AMP and some PGRP genes in response to *M. luteus* treatment and the induction levels observed for several AMP genes after *M. anisopliae* and *B. subtilis* exposures. Therefore, our findings suggest that Relish plays a critical role in inducing the humoral immune response in *Drosophila* S2 cells, contributing primarily to the response against *M. luteus* and, to a lesser extent, to the responses against *B. subtilis* and *M. anisopliae*.

## Introduction

The fruit fly is used as a model organism to study various fundamental intracellular processes (Bellen & Yamamoto, 2015), including the activation of the humoral immune response. The genetic basis of the response is highly conserved (Buchon et al., 2014). A key role in innate immunity is played by a group of κB nuclear transcription factors (NF-κB), which have originally been found in mammalian B cells (Sen & Baltimore, 1986, Hayden & Ghosh, 2011). In *Drosophila*, the NF-κB pathway primarily responds to infection by stimulating the production of antimicrobial peptides in the fat body, which is equivalent to some extent to the mammalian liver (Hetru & Hoffmann, 2009). Activation of the humoral immune response is mediated by two signaling pathways, IMD and Toll (Govind, 2008, Buchon et al., 2014) (Fig. 1). The IMD pathway is activated by Gram-negative (Gram (–)) bacteria upon direct contact of bacterial DAP-type peptidoglycans with the peptidoglycan recognition protein LC (*PGRP-LC*) (Kaneko et al., 2006, Lim et al., 2006, Westlake H., 2024). Three isoforms of the PGRP-LC receptor are known: LCx-RA, LCy-RC, and LCa-RB (Kaneko et al., 2004, Werner et al., 2003, Kleino & Silverman, 2014). *Micrococcus luteus* and *Bacillus megaterium* peptidoglycans (PGNs) bind to PGRP-LCx in vitro (Mellroth et al., 2005). However, it is widely recognized that DAP-type PGN of *B. subtilis* poorly binds to the receptor in vivo and has a minimal activating effect on IMD pathway (Gobert et al., 2003, Kaneko et al., 2005, Kaneko et al., 2004, Hedengren-Olcott et al., 2004, Ferrandon et al., 2007, Vaz et al., 2019, Westlake H., 2024).

**Figure 1.**
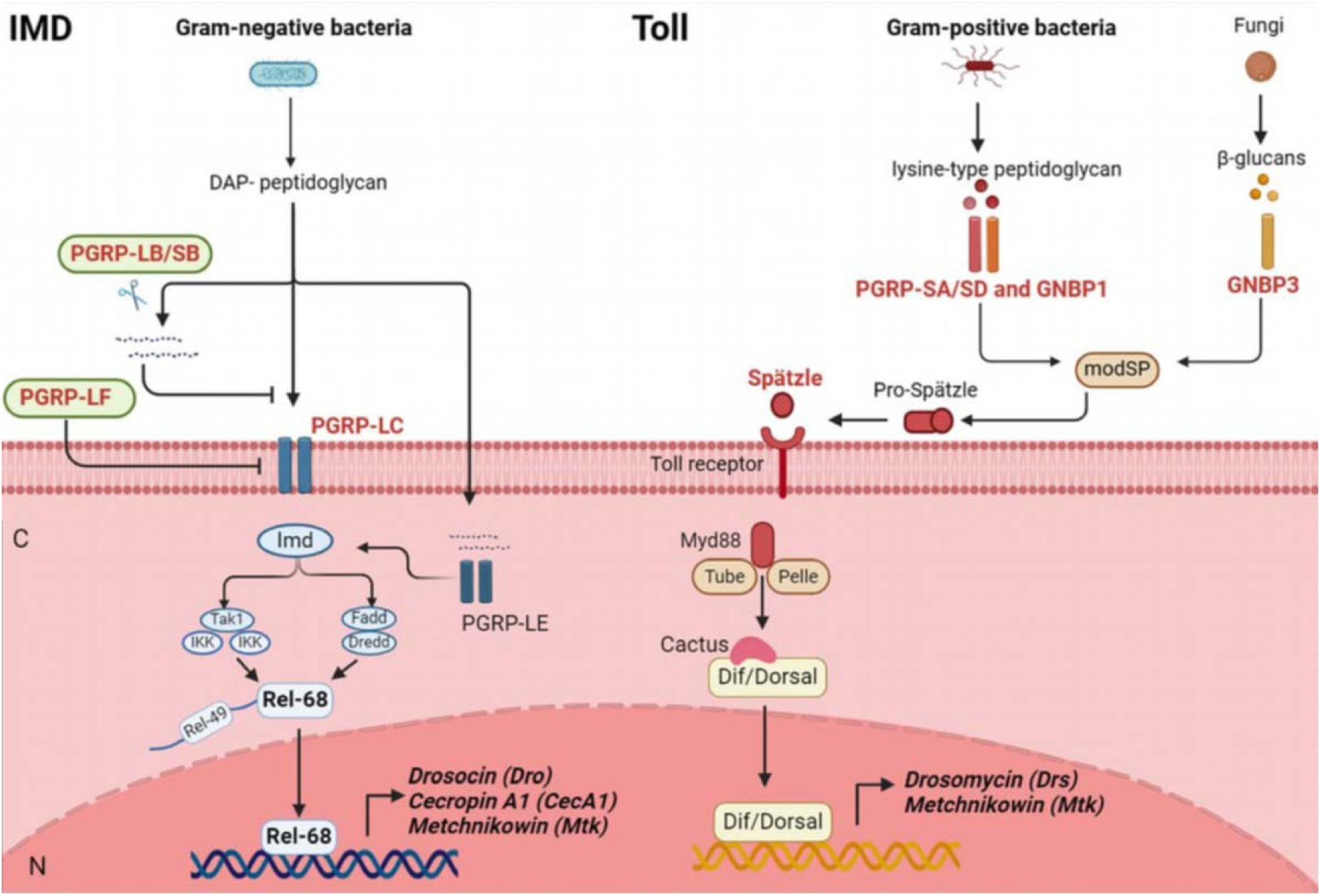
Microbe recognition and the corresponding signaling pathways induced in *D. melanogaster*. The Toll and IMD pathways are the main regulators of the humoral immune response in *D. melanogaster*. The pathways work together to orchestrate the complex immune response and thus regulate the AMPs production. The IMD pathway is activated upon the direct recognition of diaminopimelic acid (DAP)-type peptidoglycan of Gram(–) bacteria by PGRP-LC and PGRP-LE. Once activated, the IMD pathway leads to partial proteolysis and nuclear translocation of the NF[κB-family transcription factor Relish, which activates expression of AMP genes, such as *Drosocin*, *Cecropin A1*, *Attacin*, and *Metchnikowin*. In the Toll pathway, lysine-type peptidoglycans of Gram(+) bacteria are recognized by PGRP-SA and GNBP1. β-Glucans of yeasts and fungi are recognized by GNBP3. This recognition activates the proteolytic cascade that results in a culmination of the cytokine Spätzle. Spätzle binding with the Toll receptor leads to the recruitment of Myd88, Tube, and Pelle, which then trigger degradation of Cactus (IκB) to release Dorsal and/or Dif. The factors are translocated into the nucleus and activate the expression of AMP genes, such as *Drosomycin* and *Metchnikowin*.

It should be mentioned that 13 PGRPs are encoded by the *Drosophila* genome. Through alternative splicing, their genes produce more than 20 proteins, which perform various functions in the regulation of immune responses (Monahan et al., 2016). While some of them have amidase activity, others are implicated in the IMD and Toll pathways (Fig. 1) (Kurata, 2014). In contrast to the IMD pathway, the Toll signaling pathway has a more complex pathogen recognition cascade. First, a lysine-type peptidoglycan of Gram-positive (Gram (+)) bacteria interacts with the peptidoglycan recognition protein SA/SD (*PGRP-SA*/*SD*) and Gram (–) bacteria-binding proteins (Gram-negative bacteria binding protein 1 (GNBP1)). Its paralog GNBP3 interacts with fungal β-glucans(Gobert et al., 2003). These ligand-binding receptors trigger the activation of modular serine protease (modSP) (Gottar et al., 2006). Then modSP activates the Spätzle-processing enzyme (SPE), which cleaves Spätzle and eventually triggers the Toll receptor-mediated activation of the immune response (Buchon et al., 2009) (Fig. 1). The complex activation cascades of the IMD and Toll signaling pathways lead to hyperexpression of AMP genes in the fat body (Fig. 1) (Hetru & Hoffmann, 2009).

Some AMP genes are mainly regulated by the IMD signaling pathway, as is the case with *Attacin* (*Atta*), *Drosocin* (*Dro*), and *Cecropin A1* (*CecA1*), while others are predominantly regulated by the Toll pathway, such as with *Drosomycin* (*Drs*) (Lemaitre et al., 1997, Carboni et al., 2022, Chowdhury et al., 2019a). *Metchnikowin* (*Mtk*) is equally regulated by both pathways (Chowdhury et al., 2019a, Levashina et al., 1998) (Fig. 1). Recently, an entire cluster of Bomanin family genes was discovered. Unlike AMPs, genes in this family are thought to be more involved in the Toll pathway (Hanson et al., 2019). Specific transcription factors of the NF-κB family are required to activate the pathways. Activation of IMD-dependent genes requires the recruitment of the transcription factor Relish to their promoter regions, while activation of Toll-dependent genes requires the recruitment of the transcription factors Dif and/or Dorsal (Dushay et al., 1996, Ip et al., 1993, Govind, 2008, Stein et al., 1998) (Fig. 1). All three of these proteins are structurally similar and contain a Rel homology domain (RHD) (Chowdhury et al., 2019b). However, Relish, Dif, and Dorsal form different variants of homo- and heterodimers, which can dynamically cooperate to activate transcription of immune response genes after activation of the IMD and Toll pathways (Chowdhury et al., 2019b).

Despite the impressive amount of data, some issues remain controversial in the activation of IMD and Toll signaling pathways. In particular, this applies to the mechanisms of pathogen recognition by the PGRP receptors and subsequent activation of the IMD and Toll signaling pathways, which involve the transcription factors Relish and Dif/Dorsal, respectively. For example, it is not entirely clear why Gram (+) bacteria or fungi activate PGRP genes that initiate the IMD pathway. In addition, the molecular mechanisms by which Gram (+) bacteria or fungi activate IMD pathway-dependent genes remain poorly understood. Previously, several articles have suggested a likely Relish-dependent activation of the immune response in the presence of Gram (+) bacteria or fungi (Hedengren-Olcott et al., 2004). It has been shown that there is a significant reduction in *Diptericin* transcription in Relish mutants after infection with *Lactococcus lactis*, *Bacillus subtilis*, *Micrococcus luteus*, *Staphylococcus aureus*, *Metarhizium anisopliae*, etc (Hedengren-Olcott et al., 2004). It was also shown that the induction of the *CecA1* by *M. luteus* requires Relish, whereas the induction of the *CecA2* requires Dif (Hedengren-Olcott et al., 2004). However, transcription also decreases in Relish mutants in the control group after injection of Ringer’s solution to flies, which makes it difficult to assess the specificity of this effect (Hedengren-Olcott et al., 2004). More contradictory results have been obtained in fly survival experiments. For example, Hedengren et al. showed a significant contribution of Relish mutants to fly survival after infection with various fungi (Hedengren et al., 1999), although Vidal et al. later failed to confirm this (Vidal et al., 2001). Consequently, these findings fail to offer a comprehensive understanding of how Gram (+) bacteria and fungi contribute to the activation of the immune response that is dependent on Relish. Unlike flies, the S2 cell line is a more convenient model for studying the molecular mechanisms of immune response activation. For example, it is known that both pricking and injecting methods induce an injury that by itself triggers the induction of the IMD and Toll pathways in the fly (Lemaitre et al., 1997, Westlake H., 2024). Cell culture minimizes these unwanted effects, thereby enabling a more precise evaluation of how individual transcription factors or receptors influence the immune response. Finally, previous studies have used outdated methods such as Northern blot analysis, making it difficult to assess the contribution of Gram (+) bacteria and fungi to Relish-dependent activation of the immune response. We therefore studied the involvement of Relish in the activation of humoral immune responses to *M. luteus*, *B. subtilis*, and *M. anisopliae* in the S2 cell line. We have expanded the list of analyzed genes such as *Mtk*, *Dro* and some PGRP genes. Our findings demonstrate that both *E.* с*oli* and *M. luteus* strongly trigger the immune response and that Relish is recruited to the promoters of the AMP and several PGRP genes in all cases. In comparison to *M. luteus*, a less significant immune response was observed with *M. anisopliae* and *B. subtilis*. However, the recruitment of Relish and the activation of AMP genes were detected again, although their levels were lower. Our findings suggest that Relish contributes to the activation of immune responses to the Gram (+) bacteria and fungus and that its contribution is pathogen specific.

## Material and methods

### Ethics statement

Animal handling for antibody production was carried out strictly according to the procedures outlined in the NIH (USA) Guide for the Care and Use of Laboratory Animals. The protocols used were approved by the Bioethics Committee of the Institute of Gene Biology, Russian Academy of Sciences. All procedures were performed under the supervision of a licensed veterinarian under conditions that minimize pain and distress.

### S2 cell culture

S2 cells were cultured in Schneider’s *Drosophila* medium (Sigma) supplemented with 10% fetal bovine serum (HyClone) and 1×Penicillin/Streptomycin (HyClone). Cells were maintained and passaged with simple pipetting once a week and replaced with new ones from the museum (liquid nitrogen) approximately once a year.

### Antibodies

Antibodies against Relish (residues 451–628 of isoform A) were produced in our laboratory (Fig. 3A, Supplemental Fig. 2a, b). Several animals were used for immunization; the specificity of all antibodies was the same. Pre-immune normal rabbit IgG was used as a negative control.

### Protein immunoprecipitation (IP) and Western blotting (WB)

IP and WB were carried out as previously described (Kachaev et al., 2018) with some modifications. For IP, a 0-to 12-hour embryonic nuclear extract was used. The extract was treated with nucleases (1 U/μL DNase I (Thermo Scientific) and 10 μg/μL RNase (Thermo Scientific)). Approximately 2 mg of polyclonal rabbit antibodies (anti-Relish antibodies and IgG) were coupled to protein A-Sepharose (Sigma). The Sepharose pellet was washed with a buffer containing 300 mM NaCl. S2 cell lysates from control and treated cells (approximately 0.5 million cells per well) were collected, clarified from cell debris by centrifugation, and used to conduct WB. Image acquisition and quantification were performed using a ChemiDoc imaging system and ImageLab software (Bio-Rad).

### Chromatin immunoprecipitation (ChIP) assay

ChIP assays were carried out according to (Kachaev et al., 2018, Kachaev et al., 2019). S2 cells were crosslinked with 1% formaldehyde (Sigma), lysed, and centrifuged to collect the nuclear pellets. Chromatin was then segmented by sonication in a Bandelin Sonopulse 3100 HD ultrasonic homogenizer. Chromatin was immunoprecipitated using anti-Relish antibodies and IgG. The antibody amount was about 2 μg in each case. The input and immunoprecipitated fraction were analysed by RT-PCR with appropriate primers.

### Cultures of E. coli, M. luteus, and B. subtilis and M. anisopliae spores

#### Escherichia coli

BL21(DE3) cells were grown to an optical density of 0.5–0.8 and cultured aerobically in the LB medium at 37°C. The cells were centrifuged at 3000 *g* for 5 min and washed three times with 10 volumes of milliQ water (mQ-H_2_O). Finally, 10 volumes of mQ-H_2_O were added to the cell pellet. The cell culture was inactivated at 65°C for 20 min and stored at –20°C. To treat S2 cells, the *E. coli* culture was used at a 1:1000 dilution; the dose was 0.03 OD.

#### Micrococcus luteus

VKM ac2230 cells were grown to an optical density of 0.5–0.8 and cultured aerobically at 30°C. The liquid medium contained peptone, 10 g; glucose, 5 g; yeast extract, 5 g; NaCl, 5 g; distilled H_2_O, to 1000 ml; pH 7.0–7.2 (+100 µl of 10 N NaOH). The cells were washed three times with 10 volumes mQ-H_2_O. Finally, 2 volumes of mQ-H_2_O were added to the cell pellet. The cell culture was inactivated at 65 °C for 20 min and stored at –20 °C. To treat S2 cells, the *M. luteus* culture was used at a 1:50 dilution; the dose was 1.2 OD.

#### Bacillus subtilis

168 cells were grown to an optical density of 0.5–0.8 and cultured aerobically in the LB medium at 37 °C. The cells were centrifuged at 3000 *g* for 5 min and washed three times with 10 volumes of mQ-H_2_O. Finally, 2 volumes of mQ-H_2_O were added to the cell pellet. The cell culture was inactivated at 65 °C for 20 min and stored at –20 °C. The optimum activation of the immune response in S2 cells was observed when the *B. subtilis* culture was used at a 1:25 dilution and the dose was 2.4 OD.

#### Metarhizium anisopliae

VKM F-1490 spores were grown on LB agar at room temperature for one week. Spores were washed out of the Petri dish with sterile mQ-H_2_O. The *M. anisopliae* suspension was filtered through four layers of a thick sterile bandage. Fungal spores were counted in Goryaev’s chamber. The spore size is 3–4 µm. Spores were inactivated at 65°C for 20 min and stored at –20°C.

### RNA extraction and reverse transcription–quantitative PCR (RT-qPCR)

Total RNA was isolated from S2 cells using the TRIZOL reagent (MRC). Total RNA was reverse-transcribed using a Biolabmix kit (R01-50), and qPCR was performed using Taq DNA polymerase (Biolabmix) and SYBR green (Sigma). Gene expression levels were normalized to the control gene *rp49*. The relative 2^-△△CT^ method was used for data analysis. All primers are listed in Ssupplementary.

### RNA interference

RNA interference of Relish (Relish RNAi) was performed using a TranscriptAid T7 High Yield Transcription kit (Thermofisher). Chimeric primers containing T7 promoter sequences (indicated in the primer list) were synthesized to target the Relish gene. The primers were used to synthesize the first PCR product according to the manufacturer’s protocol. The product was then used as a template to synthesize the second PCR product with a primer targeting the T7 promoter sequence. The second PCR product was used for RNA synthesis. RNA synthesis was carried out at 37°C for 3 hours. After RNA synthesis, ethanol precipitation was performed, and fresh mQ-H_2_O was added. Adapter synthesis was performed by slowly decreasing the temperature from 70°C to 4°C using a 1x annealing buffer (a 10x annealing buffer contained 0.2 M Tris-HCI, pH 7.5, 0.1 M MgCI_2_, and 0.5 M NaCI). For RNAi, 20 μg of double-stranded RNA (dsRNA) targeting Relish was added to 2 million S2 cells. The resulting mixture was incubated for 5 days. The efficiency of RNAi was assessed by measuring the reduction in *Relish* gene transcription and Relish protein expression levels.

## Results

### Activation of humoral immunity in S2 cells by various pathogens

When studying the humoral immune response in insects, the expression levels of the specific pathway components are important to estimate in various tissues, organs, stages, and, of course, cell cultures. These levels show strong diversity in S2 cells (Supplemental Tab 1, adapted from Fly Base (Gramates et al., 2022)). It was of particular interest to assess the levels of transcription factors that are recruited to gene promoters after activation of innate immunity. A high level of transcription was observed for *Relish*, while expression of *Dif* and *Dorsal* was quite low (Supplemental Tab.). Treatment of S2 cells with *E. coli* is known to increase the transcription of Relish (Bonnay et al., 2014). Moreover, a high level of transcriptional activation of *Relish* has been previously found in a transcriptome analysis after infection of flies with *Staphylococcus aureus* (Troha et al., 2018). We also observed an increase in the transcription level of *Relish* after treating S2 cells with pathogens (*E. coli* and *M. luteus*), while the transcription levels of *Dif* and *Dorsal* remained almost unchanged (Supplemental Fig. 1a). The data imply that the IMD pathway is employed predominantly in S2 cells. A Toll-dependent response is expected to be very limited in S2 cells.

It is crucial to comprehend how pathogens precisely activate the Toll and IMD signaling pathways in addition to examining the role of transcription factors. *M. luteus* is among the bacteria that are widely used to activate the Toll pathway. They are very easy to cultivate, and *M. luteus* infection is not lethal to flies (Troha & Buchon, 2019). It is noteworthy that using *M. luteus* as an inducer of only the Toll pathway causes some controversy (Vaz et al., 2019, Hedengren-Olcott et al., 2004). Therefore, the fungus *M. anisopliae* was additionally used in our study, and a few experiments were carried out to compare the mechanisms of the immune response for Gram (–) vs. Gram (+) and fungal pathogens.

We compared the effects of the three pathogens *E. coli*, *M. luteus*, and *M. anisopliae* spores on the IMD- and Toll-dependent gene induction. The pathogens demonstrated the expected phenotypes on Petri dishes (Fig. 2a). The activation levels of *Drs, CecA1, Mtk,* and *Dro* were assessed after treating S2 cells with *E. coli* (control for IMD induction), *M. luteus,* and *M. anisopliae*. Consistently high levels of transcriptional activation were observed for *CecA1*, *Mtk*, and *Dro* after treating S2 cells with both *E. coli* and *M. luteus* inactivated cultures overnight (Fig. 2b, c). At the same time, the activation of *Drs* was found to be less significant and less stable after treatment with the pathogens (Fig. 2c). It is worth noting here that 20-fold higher amount of inactivated bacteria was necessary for activating the AMP genes with *M. luteus* compared to *E. coli*. Recently, it has been shown similarly that a high titer of bacteria is required to induce an immune response with *M. luteus* (Chen et al., 2022). Less significant effects on AMP gene transcription were observed after treating S2 cells with inactivated *M. anisopliae* spores for 3 hours (Fig. 2d). Unlike with *M. luteus*, even a large number of fungal spores did not activate AMP gene transcription to the same extent as bacteria. Lastly, we examined the activation states of many components (*GNBP1, GNBP3, IMD, Spätzle, Pelle, Tube,* and *Cactus*) involved in the activation cascade of the humoral immune response in our model system. Transcription levels of these genes did not exhibit any statistically significant alterations following S2 cell treatment with *E. coli*, *M. luteus*, or *M. anisopliae* (Supplemental Fig. 1b).

**Figure 2.**
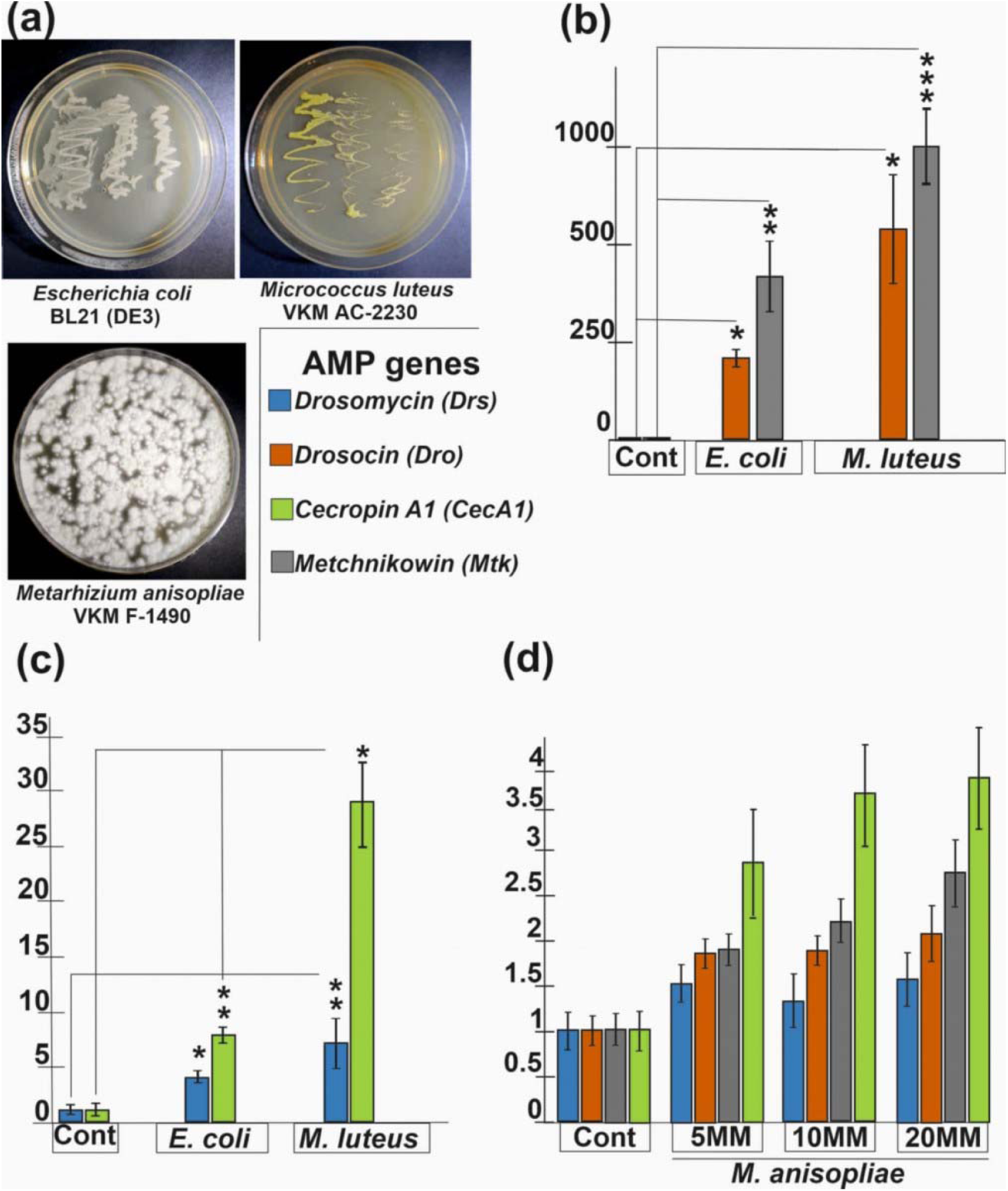
Expression levels of Drs, Dro, CecA1, and Mtk after triggering the IMD and Toll pathways in S2 cells. (a) Inoculation of *E. coli* and *M. luteus* bacteria and spores of the fungus *M. anisopliae* on Petri dishes, as well as a list of AMP genes. (b, c), and (d) AMP gene expression levels before (Cont, control) and after treatment with *E. coli* or *M. luteus*. In Figure 2c, the *CecA1* expression values in the bar graph were divided by 20. Inactivated cultures of *E. coli* and *M. luteus* were used at dilutions of 1:1000 and 1:50, respectively; *M. anisopliae* spores were added to S2 cells at 5 million (MM), 10 MM, and 20 MM. Three independent experiments were performed. Whiskers show the standard deviation. Asterisks show the statistical significance: (*) P-value ≤ 0.05, (**) P-value ≤ 0.01, or (***) P-value ≤ 0.001 as calculated in the *t*-test.

To summarize, IMD-pathway dependent genes (*CecA1* and *Dro*) predominantly showed fairly high levels of activation after treatment of S2 cells with *M. luteus*. Moreover, similar activation patterns of all AMP genes were detected after triggering the immune response with Gram (–) and Gram (+) bacteria. Significant levels of AMP gene activation were observed after 3-hour incubation of S2 cells with fungal spores. Finally, *GNBP1, GNBP3, IMD, Spätzle, Pelle, Tube,* and *Cactus* were not activated in the S2 cell model system.

### Activation of humoral immunity with M. luteus and M. anisopliae leads to the recruitment of Relish to the promoter regions of the AMP genes in S2 cells

Since the key transcription factor Relish is expressed to a sufficiently high level in S2 cells (Supplemental Fig. 1a, Tab), it is possible that Relish acts as a key regulator of immune response gene transcription in both Gram (+)- and Gram (–)-induced responses. Relish is a protein with a molecular weight of ∼110 kDa. Once the immune response is activated by Gram(–) bacteria, Relish undergoes proteolytic cleavage into two fragments with molecular weights of approximately 68 (Rel68) and 49 (Rel49) kDa (Stöven et al., 2000). The larger fragment includes the RHD and IPT domains and is imported into the nucleus and recruited to the regulatory regions of AMP genes. The smaller Relish fragment remains in the cytoplasm. The amino acid sequence that undergoes proteolysis is in region 535–552 of the Relish protein (Stoven et al., 2003). We have raised polyclonal rabbit antibodies against a Relish epitope located in region 451–628, which covers the proteolytic site (Supplemental Fig. 2b). Using WB, we found that our antibodies recognize the full-length Relish protein in S2 cells and embryonic nuclear extracts (Fig. 3a, Supplemental Fig. 2a). The larger Relish fragment Rel68 with an electrophoretic mobility of approximately 70 kDa was also detected. This means that the full-length form and its larger fragment are already present in S2 cells without any external influence by pathogens. An increase in larger Relish fragment was detected after overnight treatment of S2 cells with *E. coli* and *M. luteus* inactivated cultures (Fig. 3a). Treatment with *M. anisopliae* had an undetectable effect on Relish proteolysis by WB. Thus, both Gram (–) and Gram (+) bacteria contribute to the proteolytic cleavage of Relish in S2 cells.

**Figure 3.**
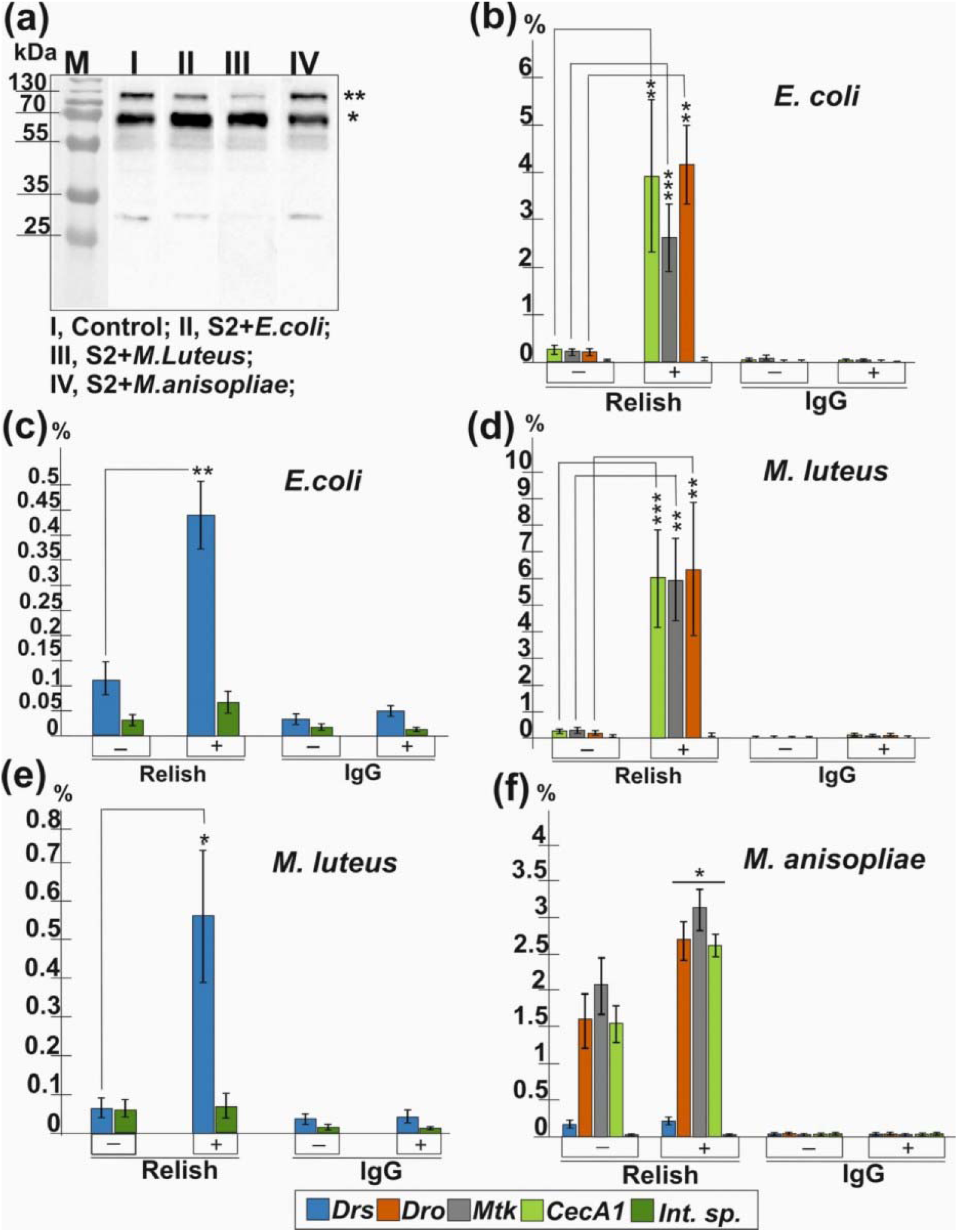
Relish recruitment to AMP gene promoters. (a) WB of S2 cell lysates with polyclonal rabbit antibodies against the Relish protein. I, Control, no treatment; II*, E. coli* treatment; III, *M. luteus* treatment; IV, *M. anisopliae* treatment. Two asterisks (**) show the full form of the Relish protein. One asterisk (*) shows the larger Relish fragment (Rel68), which forms via proteolysis. (b) Relish recruitment to the AMP genes (*CecA1, Mtk*, and *Dro*) and c) (*Drs*) upon S2 cell treatment with *E. coli*. (d) Relish recruitment to the AMP genes (*CecA1, Mtk,* and *Dro*) and e) *Drs* upon S2 cell treatment with *M. luteus*. (f) Relish recruitment to the AMP genes (*CecA1, Mtk, Dro*, and *Drs*) upon S2 cell treatment with *M. anisopliae*. Five million S2 cells were combined with *E. coli* used at 1:1000, *M. luteus* used at 1:50, or 5 MM *M. anisopliae* spores. The levels of Relish and IgG are shown as a percentage of Input (cross-linked chromatin). Three independent experiments were performed. Whiskers show the standard deviation. Asterisks show the statistical significance: (*) P-value ≤ 0.05, (**) P-value ≤ 0.01, or (***) P-value ≤ 0.001 as calculated in the *t*-test.

The above antibodies were used to analyze the recruitment of Relish to the promoter regions of the AMP genes. Relish was detected at low levels on the promoters of *Drs*, *CecA1*, *Mtk*, and *Dro* before treatment; its levels significantly increased after treatment with the bacterial pathogens (Fig. 3b-e). Although regarded as an activator of the Toll pathway, *M. luteus* induced Relish recruitment to the IMD pathway-dependent genes *Dro* and *CecA1* (Fig. 3d). Moreover, comparable levels of Relish recruitment to the AMP gene promoters were observed in S2 cells treated with *E. coli* or *M. luteus* (Fig. 3b-e). Therefore, regardless of the activation of the IMD and Toll pathways in a S2 cell culture, the Relish transcription factor plays a predominant role in the regulation of AMP gene transcription. Relish recruitment to the *Drs* promoter was noticeably lower than to the promoters of the other AMP genes (Fig. 3c, e). This apparently explains the poor activation of the *Drs* gene in S2 cells (Fig. 1c). The recruitment of Relish to the AMP gene promoters was additionally studied in S2 cells treated with fungal spores. A slight increase in Relish recruitment to *CecA1*, *Mtk*, and *Dro* was observed (Fig. 3f). Thus, although undetected by WB, Relish was activated in S2 cells by fungal spores according to the ChIP data.

Our data showed quite similar patterns of proteolysis and recruitment of the Relish transcription factor to the promoters of the AMP genes for S2 cells treated with bacteria. *M. anisopliae* had a similar but lower effect on the recruitment of Relish to the AMP genes.

### Relish RNA knockdown leads to repression of AMP gene transcription after treatment of S2 cells with M. luteus and M. anisopliae

A Relish RNA knockdown was tested for effect on transcription of the corresponding genes. Using *E. coli* and *M. luteus*, we examined the effect of the Relish knockdown on the activation of the AMP genes subsequent to the induction of the immune response. Our data showed a noticeable decrease in Relish at both the RNA and protein levels (Fig. 4a, b). As expected, the level of AMP gene transcription decreased significantly after activation of the IMD pathway with *E. coli* (Fig. 4b). Surprisingly, similar effects were observed after activating the Toll pathway with *M. luteus* (Fig. 4b). The transcription levels of all genes studied decreased significantly.

**Figure 4.**
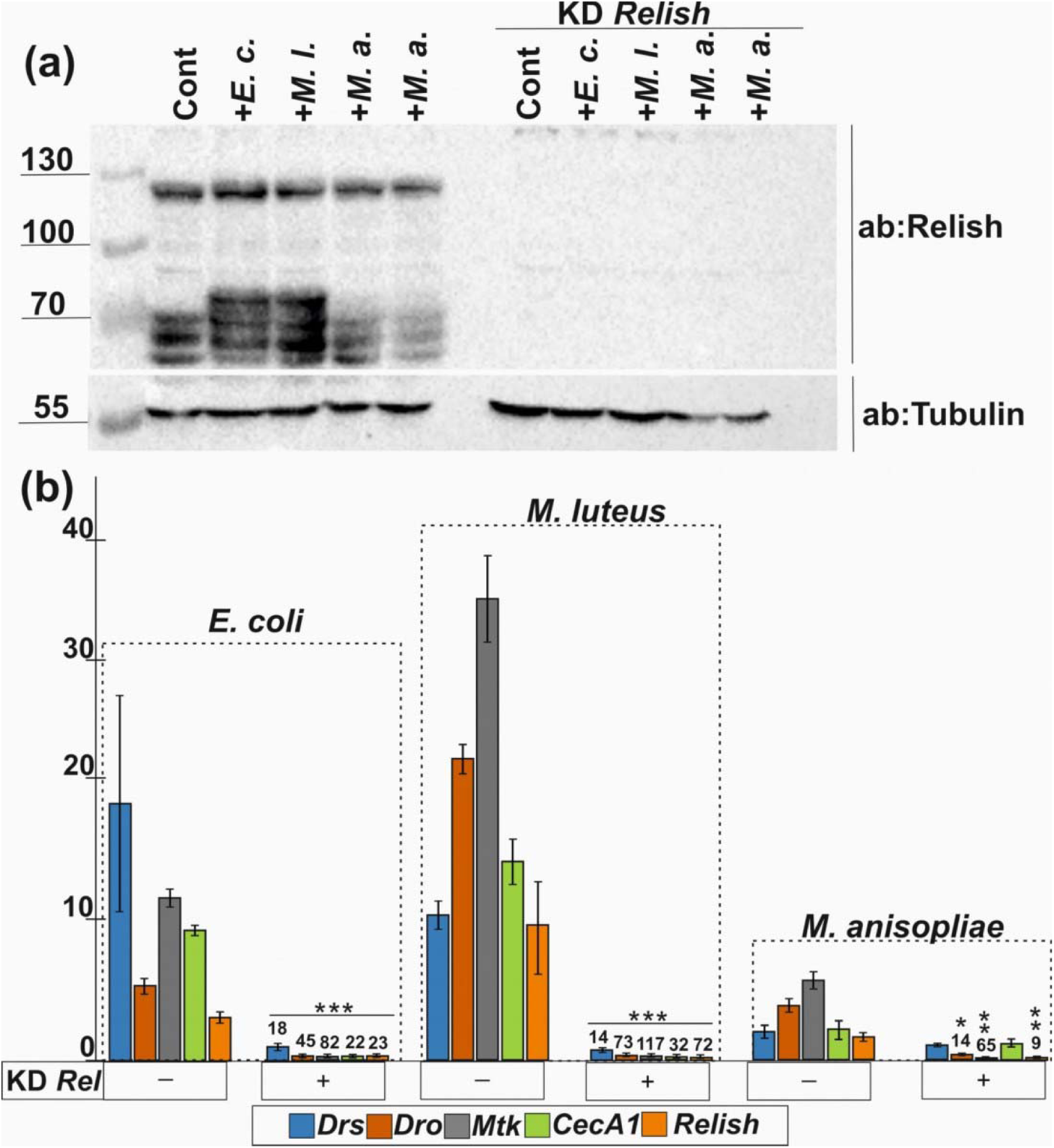
Relish RNA knockdown and activation of the immune response by *E. coli*, *M. luteus*, and *M. anisopliae*. (a) A knockdown of Relish and WB of a total protein extract from S2 cells with anti-Relish antibodies before (Cont) and after treatment with *E. coli* (*E. c.*), *M. luteus* (*M. l.*), and a double amount of *M. anisopliae* spores (*M. a.*). The full-length form of the Relish protein is detected at about 110 kDa; the proteolytic (nuclear) form is detected at about 70 kDa. A Relish knockdown in S2 cells specifically decreased the intensities of both bands (the full-length and nuclear forms) detected with Relish antibodies. Tubulin was used as a loading control. (b) Effect of a Relish RNA knockdown on transcription of the AMP genes. The activation levels of *Mtk* and *Dro* were divided by 20 in the case of bacterial treatment. Fold decreases in AMP gene transcription after the Relish knockdown are shown in Arabic numerals over the bars. Three independent experiments were performed. Whiskers show the standard deviation. Asterisks show the statistical significance: (*) P-value ≤ 0.05, (**) P-value ≤ 0.01, or (***) P-value ≤ 0.001 as calculated in the *t*-test; comparisons were performed with the values observed after activation with *E. coli*, *M. luteus*, and *M. anisopliae*.

**Figure 5.**
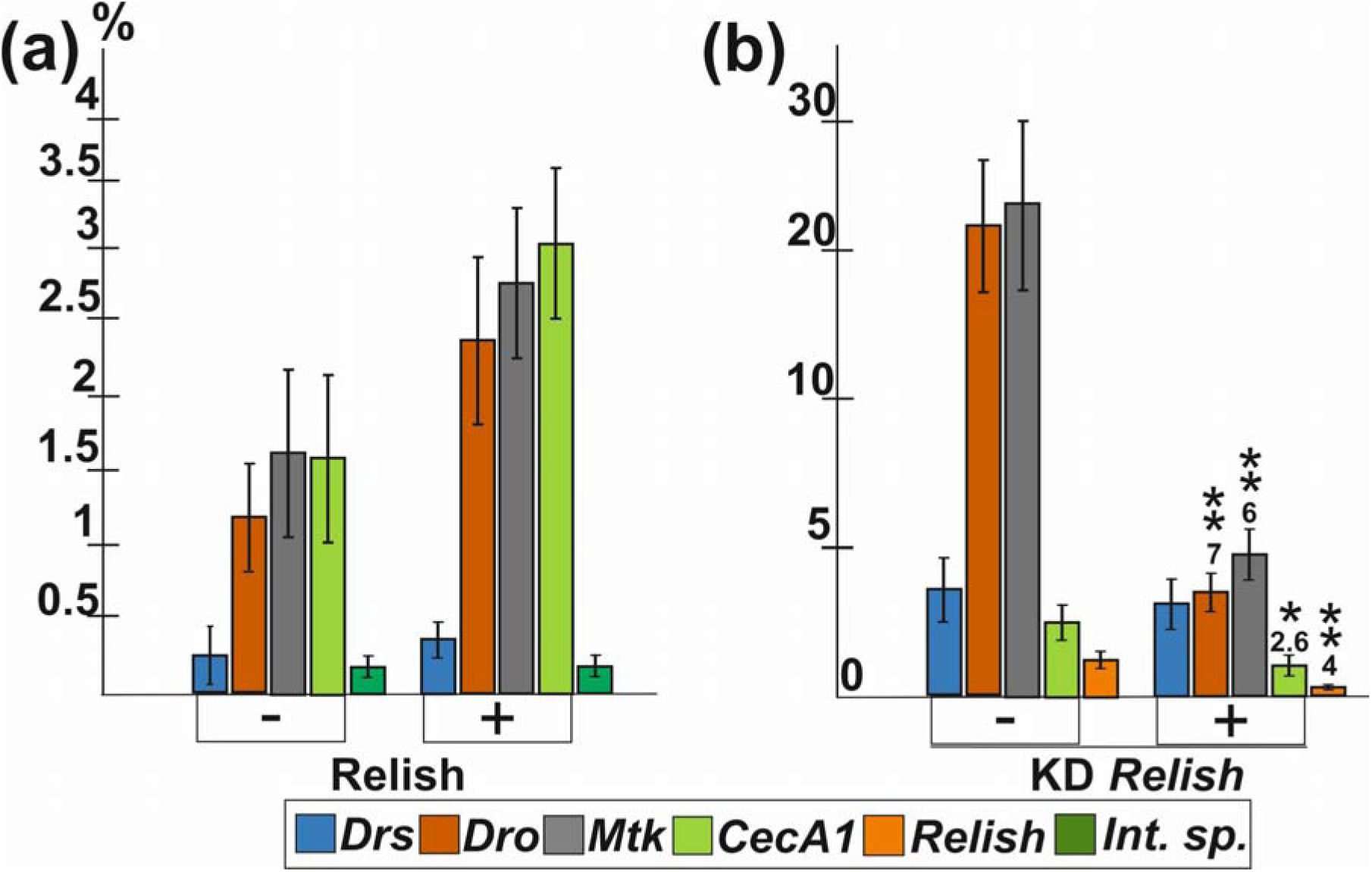
Analysis of the Relish-mediated activation of the immune response upon *B. subtilis* treatment of S2 cells. (a) Relish recruitment to the AMP genes in S2 cells treated with *B. subtilis*. ([) before treatment, (+) after treatment with *B. subtilis*. (b) Effect of a Relish RNA knockdown (KD *Relish*) on the transcription of the AMP genes in cells treated with *B. subtilis*. Fold decreases in AMP gene transcription after the Relish knockdown are shown in Arabic numerals over the bars. Three independent experiments were performed with *Relish* KD cells (b). *B. subtilis* was added to 5 million S2 cells at a 1:25 dilution. Whiskers show the standard deviation. Asterisks show the statistical significance: (*) P-value ≤ 0.05, (**) P-value ≤ 0.01, or (***) P-value ≤ 0.001 as calculated in the *t*-test; comparisons were performed with the values observed after activation with *B. subtilis*.

Since lower levels of AMP gene activation were observed after activation of the humoral immune response by treating S2 cells with *M. anisopliae*, we decided to analyze the effect of a Relish RNA knockdown on the activation of the AMP genes. Our data show that the Relish knockdown had virtually no effect on *Drs* and *CecA1* transcription, whereas *Mtk* and *Dro* transcription levels decreased significantly.

Thus, we were the first to discover the recruitment of the Relish factor to AMP gene promoters after treatment of S2 cells with inactivated *M. luteus* cultures. It is possible that this effect is associated specifically with the pathogen type or that the IMD pathway-dependent activation of the humoral immune response is predominantly active in S2 cells.

*Bacillus subtilis* shows Relish-dependent immune response activation in S2 cells

As we found high levels of activation of the AMP genes after *M. luteus*, which was Relish-dependent, we decided to analyze if another Gram (+) bacterium, *B. subtilis*, can trigger the immune response in S2 cells. We found that *B. subtilis* stimulates the expression of several AMP genes (Supplemental Fig. 3a). Their induction was less than with *M. luteus*, although twice as much bacterial were added to S2 the cells in the case of *B. subtilis*. A noticeable increase in processed Rel68 levels was additionally observed after *B. subtilis* treatment of S2 cells (Supplemental Fig. 3b), and the recruitment of Rel68 to the promoter regions of AMP genes was detected (Fig. 6a). Finally, a Relish RNA knockdown affected the induction of *Dro, Mtk* and, *CecA1*, while *Drs* activation was mostly unaffected (Fig. 6b).

**Figure 6.**
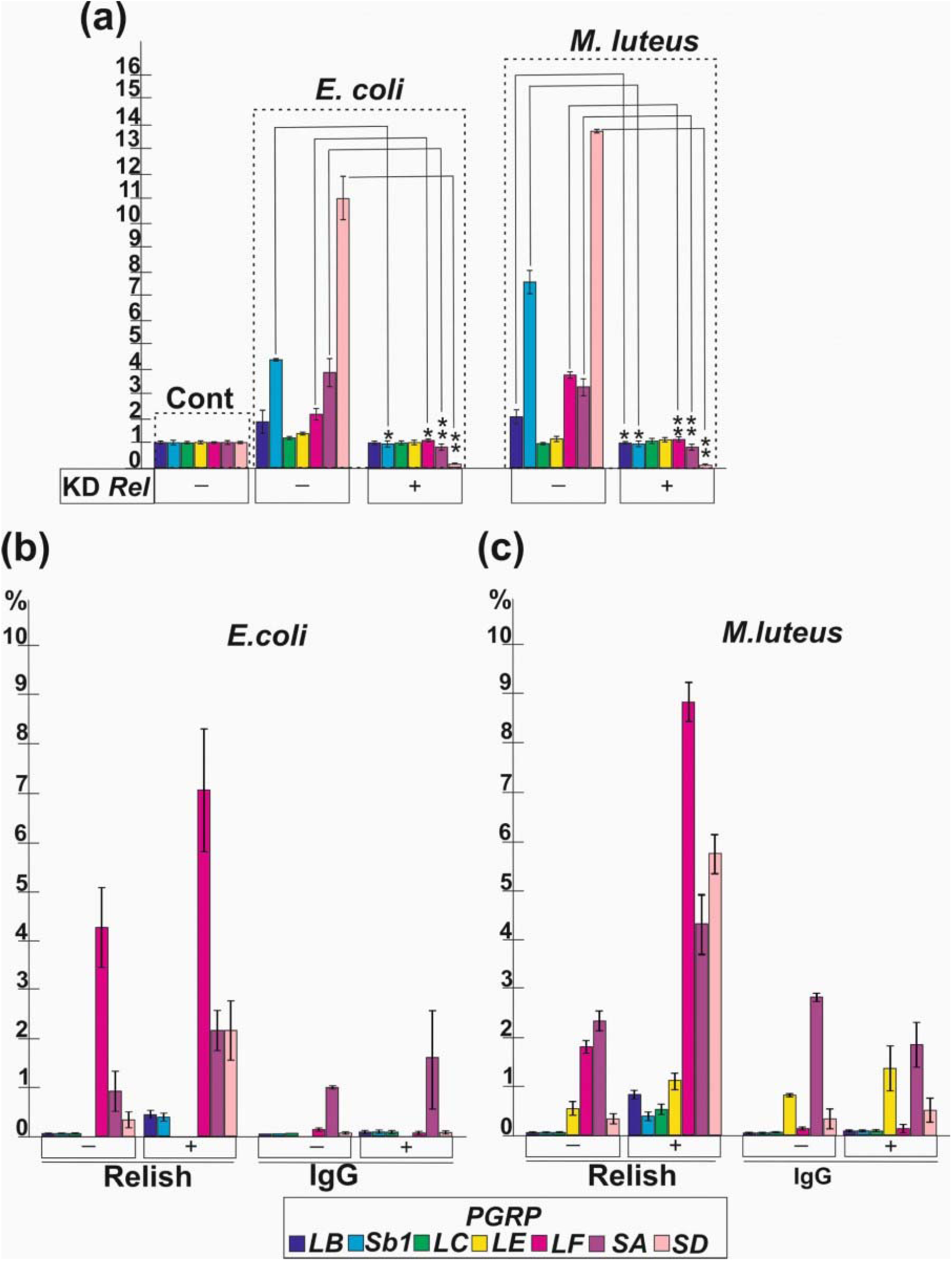
Analysis of PGRP activation and Relish recruitment in S2 cell cultures treated with *E. coli* or *M. luteus*. (a) The levels of PGRP gene activation in S2 cells before (Cont) and after *E. coli* and *M. luteus* treatment. Transcription of some PGRP genes significantly decreased after a Relish knockdown (KD). (b, c) Relish recruitment to the PGRP genes upon S2 cell treatment with *E. coli* and *M. luteus*. Three independent experiments were performed. Whiskers show the standard deviation. Asterisks show the statistical significance: (*) P-value ≤ 0.05, (**) P-value ≤ 0.01, or (***) P-value ≤ 0.001 as calculated in the *t*-test.

Thus, we confirmed that Relish contributes to the response to *B. subtilis*, although its contribution was less prominent than to the responses to the other bacteria studied. The contribution was similar to that observed in the response to fungal spores.

### Relish-dependent activation of PGRP receptors after activation of the immune response in S2 cells by E. coli and M. luteus

It has previously been shown that *E. coli* and *M. luteus* lead to the activation of some PGRP receptors in adult flies (Troha et al., 2018). We analysed the transcription activation levels of several PGRP receptor genes, including *LB, Sb1, LC, LE, LF, SA,* and *SD*. The PGRP receptors LC, LE, and LF are involved in the IMD pathway, whereas SA is involved in the Toll pathway. The participation of PGRP-SD in the Toll or IMD pathway remains controversial (De Gregorio et al., 2002, Iatsenko et al., 2016). It is possible that this receptor participates in both the Toll and IMD pathways. Amidase activity, which is inherent in the LB and Sb1 receptors, catalyzes the hydrolytic cleavage of PGNs from the respective pathogens, and it has been shown that the receptors are involved in the IMD pathway (Mellroth & Steiner, 2006, Zaidman-Rémy et al., 2006, Zaidman-Rémy et al., 2011).

We found that *PGRP-SD*, *SA, Sb1, LB,* and *LF* were activated after treating S2 cells with *E. coli* and *M. luteus*. At the same time, the activation levels of *PGRP-Sb1* and *LF* were slightly higher when S2 cells were treated with *M. luteus* (Fig. 6a). Using ChIP, we found that Relish was recruited to the promoter regions of the *PGRP-SD*, *Sb1*, *LC*, *LB,* and *LF* genes after treatment of S2 cells with *M. luteus* and *E. coli* (Fig. 6b, c). Along with the fairly high activation of *PGRP-SD* after immune response activation with *E. coli* and *M. luteus*, a significant presence of Relish was additionally observed on the promoter of this receptor gene.

Finally, we tested the effect of Relish depletion on PGRP gene transcription after activation of the immune response with *E. coli* and *M. luteus*. A statistically significant decrease in the transcription levels of *PGRP-SD, LF,* and *Sb1* was observed after activation of the immune response with either bacterium (Fig. 6a). The finding is in line with the induction of these genes and Relish recruitment onto their promoters upon pathogen treatment.

Thus, the group of the *PGRP-SD*, *LF,* and *Sb1* genes is regulated by Relish after activation by pathogens. The respective receptors are predominately related to the IMD pathway.

## Discussion

Different classes of pathogens trigger the activation of the humoral immune response. PGRPs, IMD- and Spz-type receptors, and the transcription factors Relish, Dif, and Dorsal are important for the differentiation of the humoral immune response with respect to the IMD and Toll signaling pathways. These are important markers that show the activation of a particular pathway. Finally, as noted above, AMPs genes have been classified into IMD- and Toll-dependent genes. For example, activation of *Drs* is attributed to the Toll pathway, while activation of *Dro* and *CecA1*is attributed to the IMD pathway (Lemaitre et al., 1997, Carboni et al., 2022, Chowdhury et al., 2019a). *Mtk* is expressed upon the activation of both Toll and IMD signaling pathways (Chowdhury et al., 2019a, Levashina et al., 1998).

Contrary to the above model, we found similar patterns of transcriptional activation for some AMP and PGRP genes after triggering the immune response with *E. coli* and *M. luteus*. Similar activation patterns of these genes have previously been demonstrated in a transcriptome analysis (Troha et al., 2018). Since Relish is predominantly expressed in S2 cells, we suggested that Relish may serve as a key transcription factor that regulates the activation after triggering the humoral immune response with *E. coli* and *M. luteus*. The Relish variant resulting from proteolysis (Rel68) was detected in S2 cells not only after treatment with *E. coli* but also after treatment with *M. luteus*. Proteolysis indicates the formation of an active form of Relish, which is necessary for the activation of AMP gene transcription. Indeed, it turned out that Relish is recruited to the AMP and PGRP genes to a noticeable level after activation of the immune response with *E. coli* and *M. luteus*. Moreover, a Relish RNA knockdown after treatment of S2 cells with *E. coli* and *M. luteus* led to a significant decrease in *Drs*, *CecA1*, *Mtk*, and *Dro* transcription and activation of some PGRPs. At the same time, S2 cells treated with *B. subtilis* and *M. anisopliae* showed less pronounced AMP genes activation. We did not observe an increase in the Rel68 signal after inducing an immune response in S2 cells with fungal spores, while *B. subtilis* caused a more pronounced signal increase. In the case of fungal induction, a Relish RNA knockdown similarly reduced transcription of *Mtk* and *Dro*. Meanwhile, KD *Relish* did not affect *Drs* transcription after treatment of S2 cells with *B. subtilis* and *M. anisopliae*. Thus, all four pathogens differentially activate immune response genes in S2 cells.

It has previously been shown that Relish may be a key factor in the activation of the humoral immune response (Hedengren-Olcott et al., 2004, Hedengren et al., 1999). It has been assumed that activation of the humoral response depends more specifically on the pathogen type, i.e., Gram (+) bacteria, Gram (–) bacteria, and fungi may be more or less involved in the activation of both the IMD and Toll pathways. A large set of Gram(+) and Gram (–) bacteria and fungi, including *M. luteus, B. subtilis* and *M. anisopliae,* has previously been tested, and Relish mutations have been found to significantly decrease the levels *Dpt* transcription (Hedengren-Olcott et al., 2004). Surprisingly, in adult flies, these AMPs are activated at very high levels after exposure to bacteria and fungi, whereas we did not find this to be the case in the S2 cell line. Some AMP genes studied in adult flies were also analysed in S2, such as *CecA1* and *Drs*. Our findings indicate that knockdown of Relish results in a marked reduction of transcription for both genes when *M. luteus* is present, but in adult flies, only *CecA1* experiences a decrease (Hedengren-Olcott et al., 2004). In contrast, after *B. subtilis* and *M. anisopliae*, similar effects are observed for *CecA1* in S2 cells and flies. Thus, in S2 cells and flies, a pathogen-specific activation of the immune response is observed. PGRP receptors form a complex repertoire of interactions with each other and with pathogens to ensure a rapid immune response to external stimuli (Kurata, 2014). Relish has been shown to regulate transcription of *PGRP-SD* after activation of the immune response with a mixture of *E. coli* and *M. luteus* bacteria (De Gregorio et al., 2002, Iatsenko et al., 2016). Thus, it was not entirely clear whether Relish regulates *PGRP-SD* transcription after infection solely with *M. luteus*. We found that Relish regulates *PGRP-SD* transcription after activation with *E. coli* as well as with *M. luteus*. Using ChIP, we found high levels of Relish recruitment to the promoter region of this gene. Our data indicate that Relish is directly involved in regulating *PGRP-SD, LF*, and *Sb1* expression after activation of the immune response with *E. coli* and *M. luteus*. It was surprising that transcriptional activation of *PGRP-LB*, *Sb1* (have amidase activity), and *LF* was observed after S2 cells treatment with *M. luteus*, although these receptors are involved in the IMD pathway (Persson et al., 2007, Mellroth & Steiner, 2006, Zaidman-Rémy et al., 2011, Zaidman-Rémy et al., 2006). The same studies have shown that the receptors are involved in the IMD pathway upon contact with DAP-type PGNs. Nonspecific activation of certain components of the signaling pathways has recently been observed in S2 cells treated with Gram (–) and Gram (+) bacteria. Thus, the Spätzle protein can be activated in the presence of the Gram (–) bacteria *Pseudomonas entomophila* and *Providencia rettgeri* in flies (Troha et al., 2018). These bacteria also lead to hyperactivation of the PGRP-SA receptor (the Toll pathway) (Troha et al., 2018). After treating S2 cells with *M. luteus*, we did not find transcriptional activation of PGRP-LC, which is the primary receptor of the IMD pathway. It is noteworthy that prior research has indicated a potential role of PGRP-LC in triggering the humoral immune response in mbn-2 cells and adult flies following treatment with *M. luteus* and *B. subtilis* (Choe et al., 2002). At the same time, the data have not been confirmed in independent research (Leulier et al., 2003, Kaneko et al., 2005, Kaneko et al., 2004). However, it is assumed that the PGN of *B. subtilis* may interact weakly with PGRP-LCx and slightly activate the IMD pathway (Kaneko et al., 2004, Ferrandon et al., 2007, Vaz et al., 2019, Westlake H., 2024). The reduced potency of *B. subtilis* PGN may be due to amidation of DAP residues, which are abundant in PGN from this microbe (Atrih et al., 1999, Kaneko et al., 2004). It has recently been demonstrated that the presence of WTAs (wall teichoic acids) in PGNs of Gram (+) bacteria significantly reduces their ability to bind to PGRP-LC (Vaz et al., 2019). Thus, it remains relevant to study the mechanisms whereby PGNs of Gram (+) bacteria are recognized by PGRPs and the immune response is subsequently activated via a Relish-dependent mechanism. It is important to highlight that emerging evidence points to the existence of alternative pathways for immune response activation in S2 cells, with Relish playing a role in these processes. Relish can also undergo alternative activation by cGAS–STING pathway (Goto et al., 2018). This mode merges with the canonical Imd pathway at the level of IKKβ and does not involve upstream components of the pathway such as Imd. cGAS-STING regulates a set of STING-regulated genes (Srgs) independent of PGRP-LC-Imd-Relish target genes in the fat body (Goto et al., 2018, Westlake H., 2024). Future studies are required to better characterize alternative modes of Imd pathway.

Another potential source of contradictions between the above studies may lie in the use of different strains of the same bacteria, chemically purified PGN, and types of bacteria that have a unique PGN structure. In addition, live and inactivated bacterial cultures have different effects on the immune response (Troha et al., 2018). A separate comment on the unrepresentativeness of using *M. luteus* as an inducer of the Toll pathway only has been provided by Petros Ligoxygakis’ team (Vaz et al., 2019). They have noted that *M. luteus* PGN differs from PGNs of other Gram (+) bacteria. The PGN of *M. luteus* belongs to group A2 (in which crosslinking is made by a single polymerized stem peptide), which is uncommon among bacteria (Vaz et al., 2019, Schleifer & Kandler, 1972). It has therefore been noted that these characteristics make *M. luteus* PGN not a representative choice for a Gram (+) bacterium (Vaz et al., 2019). Perhaps this explains some of our observations and, in particular, the finding that treatment of S2 cells with *M. luteus* leads to Relish-mediated activation of the AMP and PGRP genes.

To conclude, like *E. coli, M. luteus* significantly activates the humoral immune response in S2 cells by a Relish-dependent mechanism, while *M. anisopliae* and *B. subtilis* also use this mechanism, but in a modest way (Supplemental Tab 2). Relish therefore plays an orchestrating role to control the response of the immune system against various types of pathogens, including Gram (+) and Gram (–) bacteria. Our data suggest that the Relish factor plays a dominant role in transcriptional activation of immune response genes in the S2 cell line, regardless of the pathogen type. However, it is still relevant to study the contributions of all three NF-κB factors to the immune response in cells of different origin.

## Author contributions

Z.M.K., conceptualization, writing and editing of the original draft, experimental work, supervision and funding acquisition; M.G., experimental work and editing of the original draft; A.A.M., experimental work; A.V.S., experimental work; I.Y.T., editing original draft Y.A.P., editing of the original draft; N.G.S., editing original draft; V.K.C., editing original draft; Y.V.S., conceptualization and editing of the original draft.

## Acknowledgements

We thank Dr. Svetlana M. Ozerskaya (Scriabin Institute of Biochemistry and Physiology of Microorganisms, RAS, Moscow region, Pushchino, Russia) for kindly providing the strains *M. luteus* VKM Ас-2230 and *M. anisopliae* VКМ F-1490.

We thank Dr. Alexey Kulikovsky (Laboratory of Molecular Genetics of Microorganisms, Institute of Gene Biology, RAS, Moscow, Russia) for kindly providing the strain *Bacillus subtilis* 168.

We thank the Center for Precision Genome Editing and Genetic Technologies for Biomedicine (Institute of Gene Biology, RAS) for assistance with gene expression analyses.

## Data availability

The datasets analysed during the current study are available from the corresponding authors on reasonable request.

## Supplementary materials

**Supplementary Table 1.**
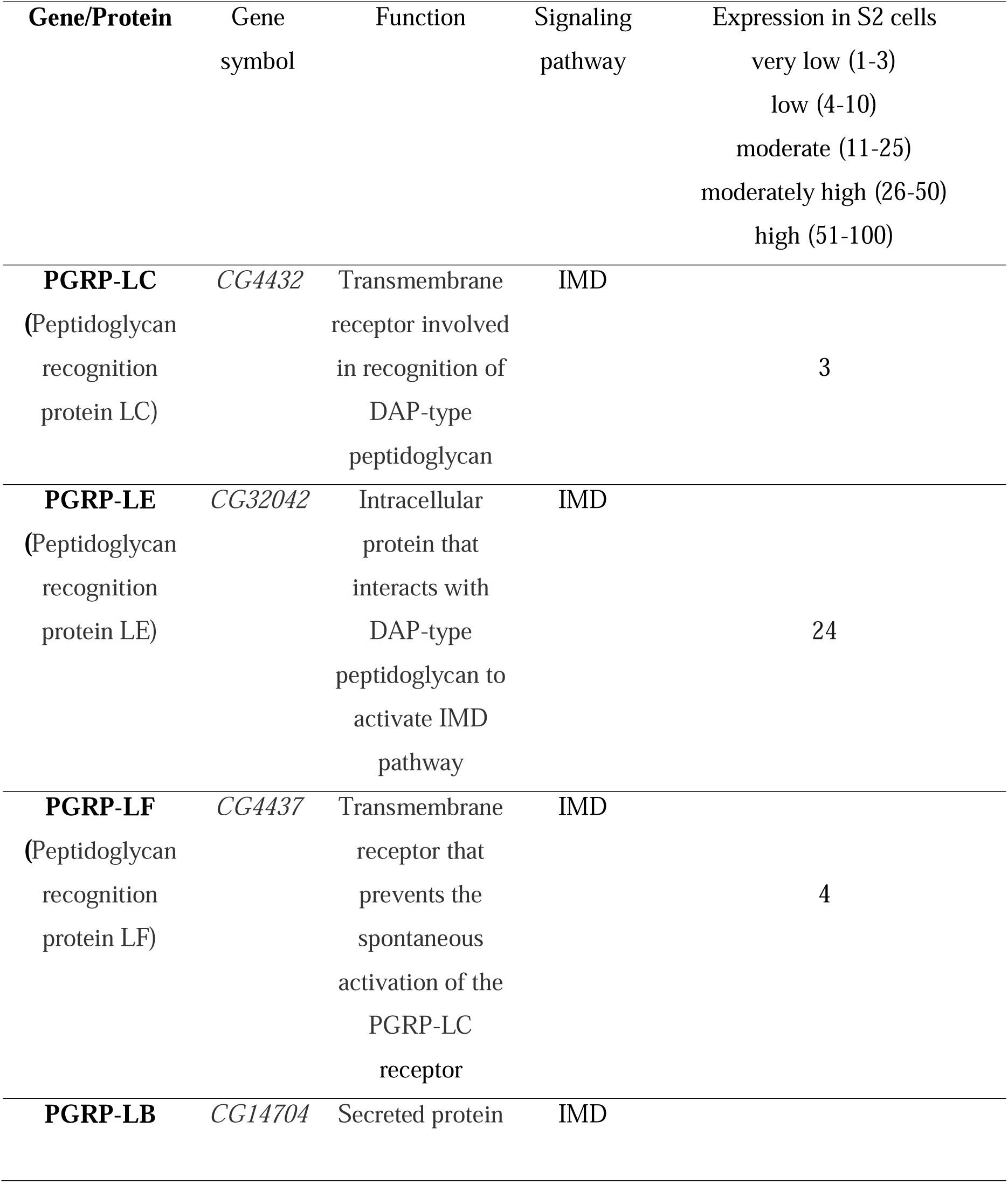
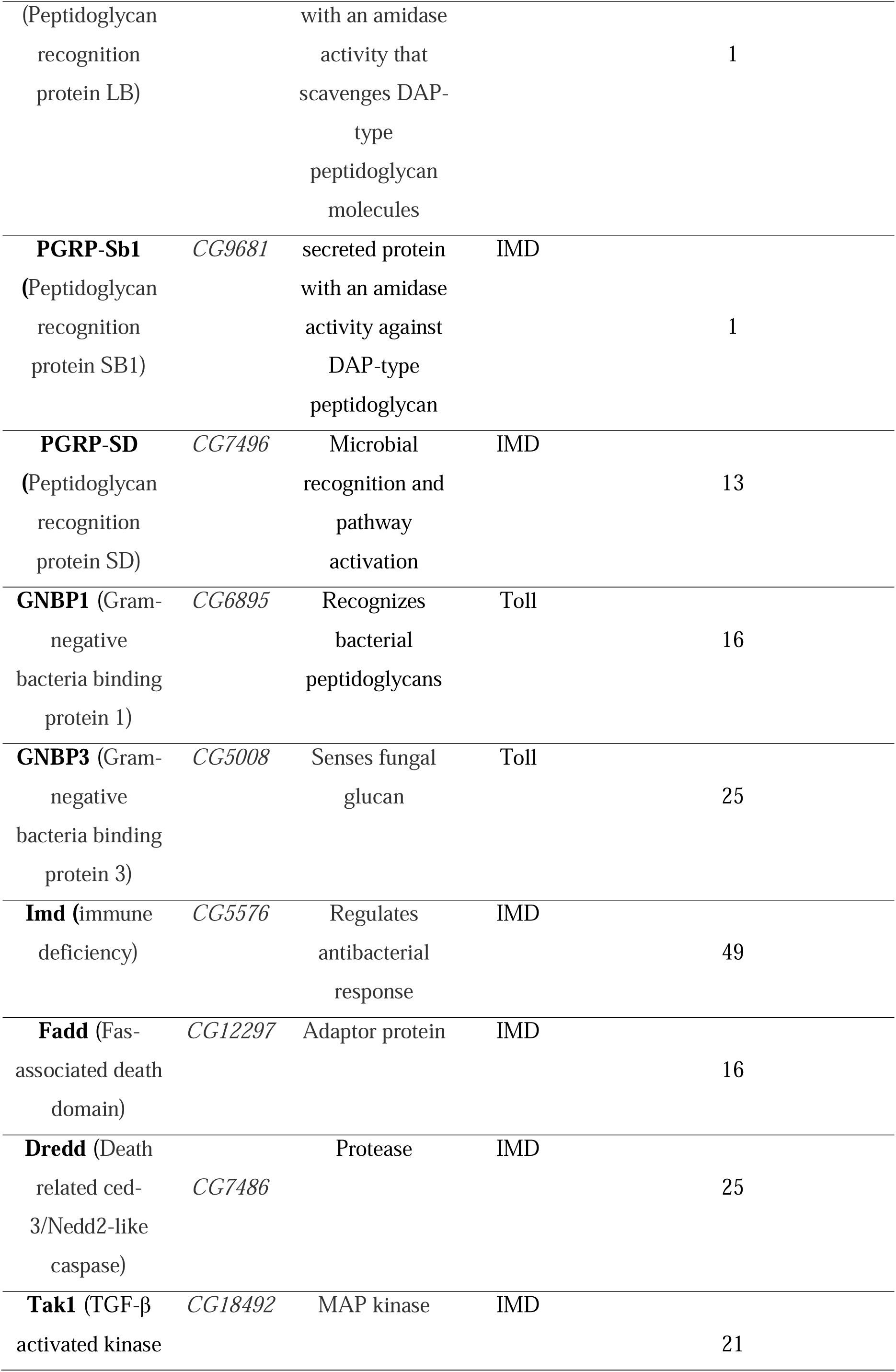
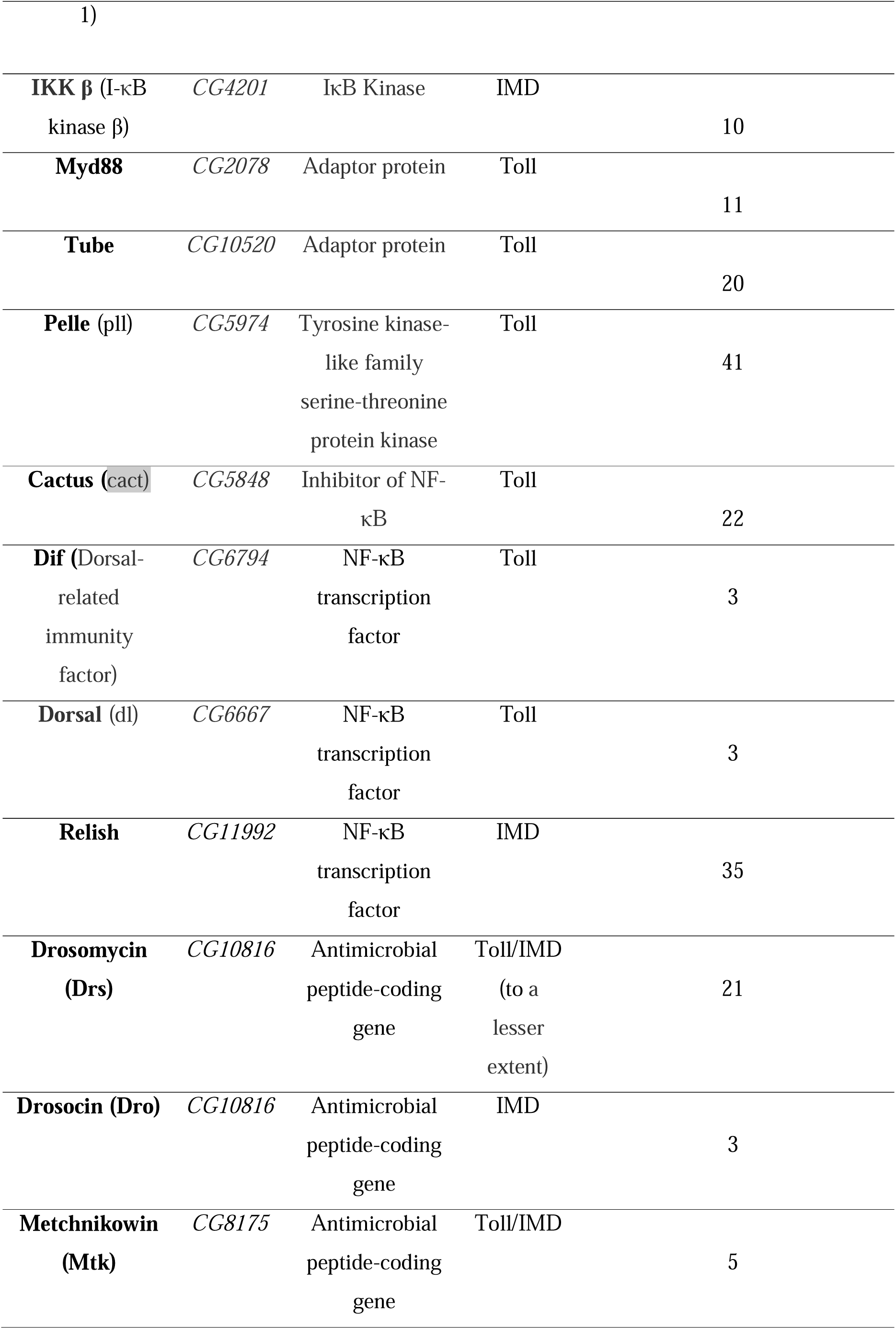
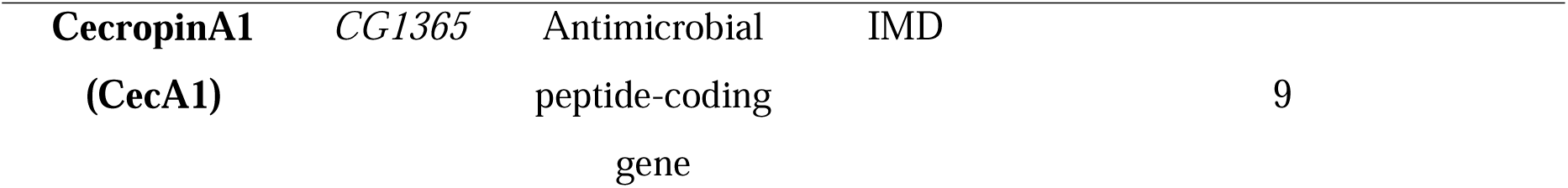
Toll and IMD main components and their expression levels in S2 cells. Adapted from Flybase (Gramates et al., 2022).

**Supplementary Figure 1.**
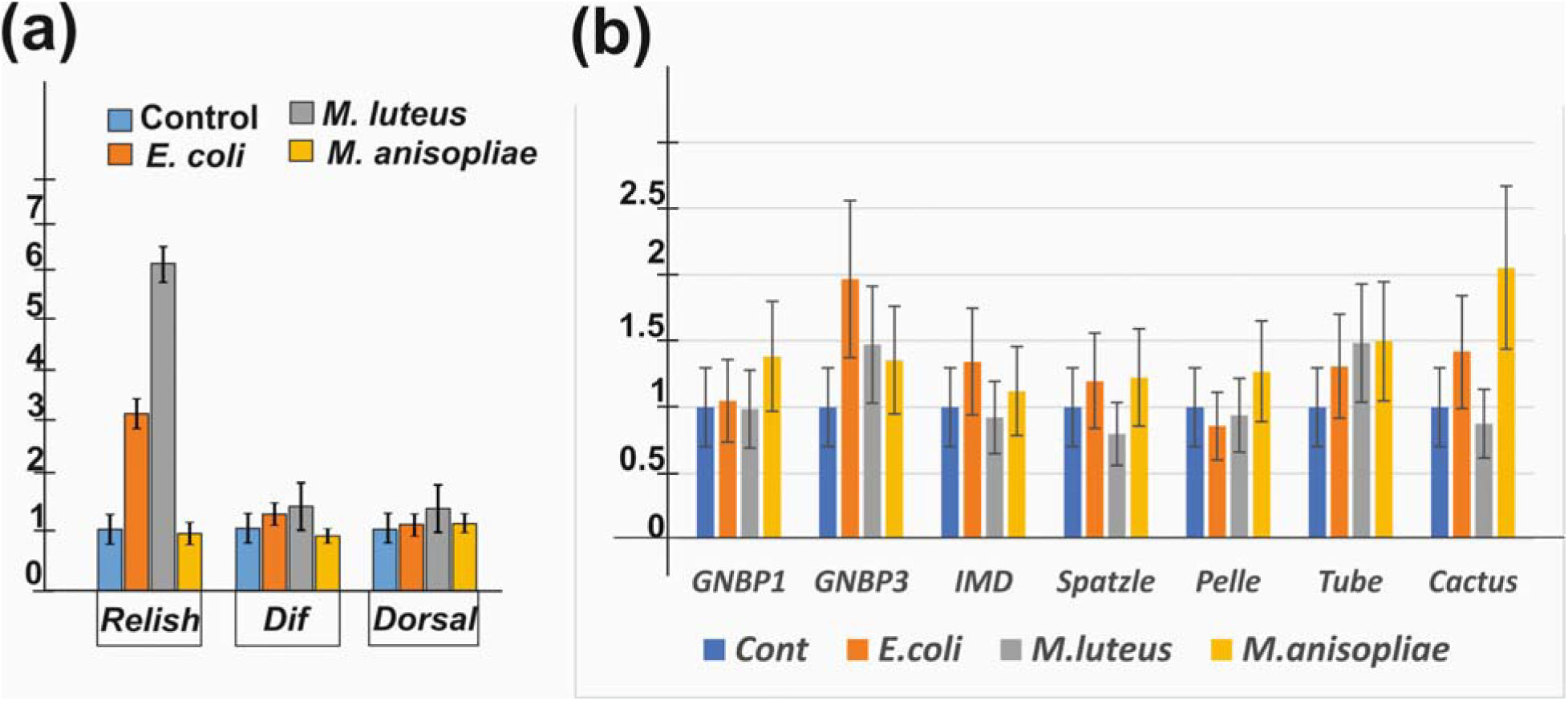
Activation of humoral immune response with *E. coli*, *M. luteus* and *M. anisopliae*. a), b) Expression levels of *Relish, Dif, Dorsal*, *GNBP1, GNBP2, GNBP3*, *IMD, Spatzle, Pelle, Tube and Cactus* after triggering immune response in S2 cells with pathogens. Three independent experiments were performed, and standard deviations are shown as error bars.

**Supplementary Figure 2.**
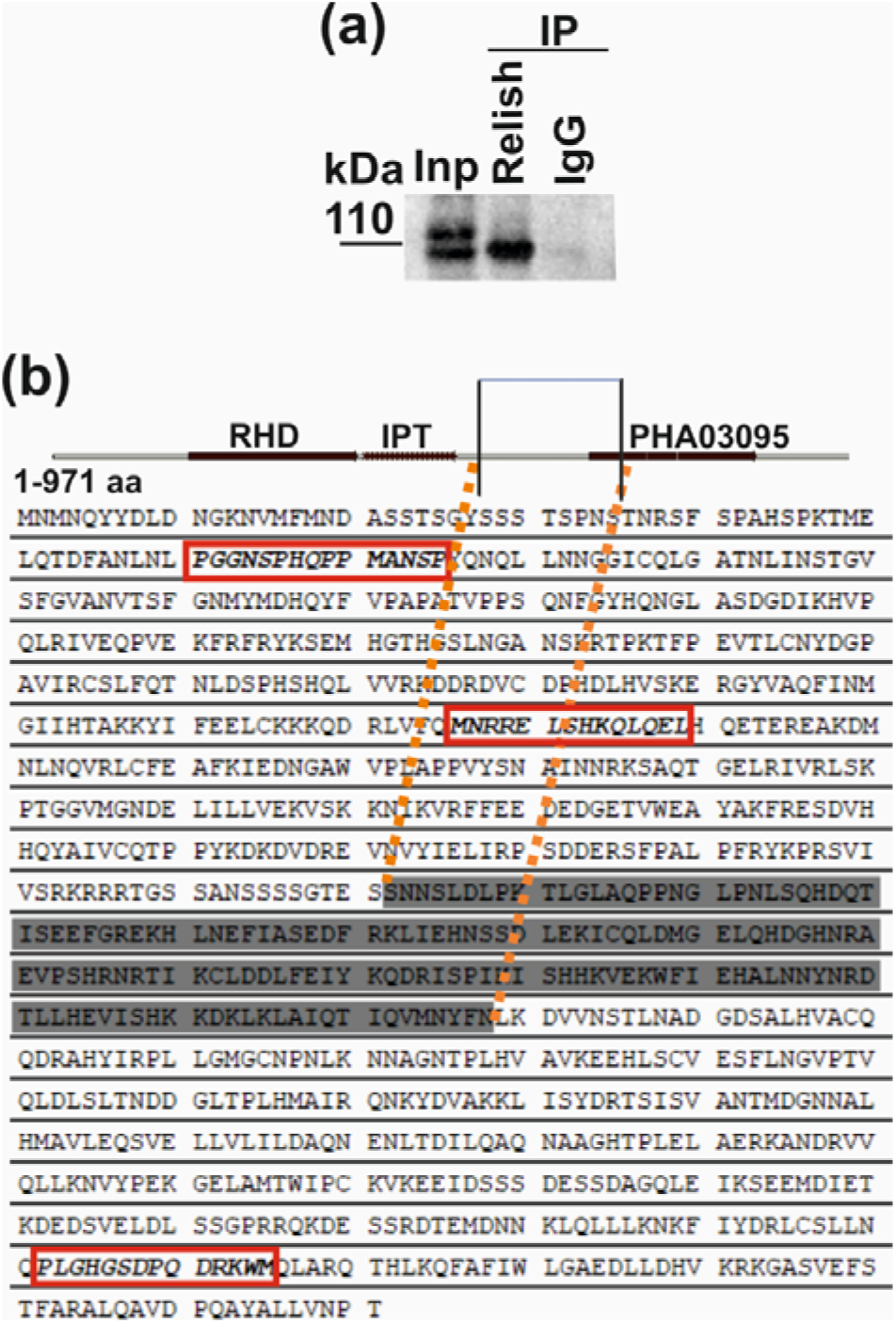
Anti-Relish antibodies. a) Immunoprecipitation (IP) and subsequent Western blot analysis detect predominantly the full-length form of the Relish protein at 110 kDa. IP was performed with a 12-h embryonic extract. b) Relish amino acids sequence. The amino acid sequence that undergoes proteolysis is in region 535–552 of the Relish protein. Framed sequences have been identified as proteolysis sites (Svenja et al., 2000).

**Supplementary Figure 3.**
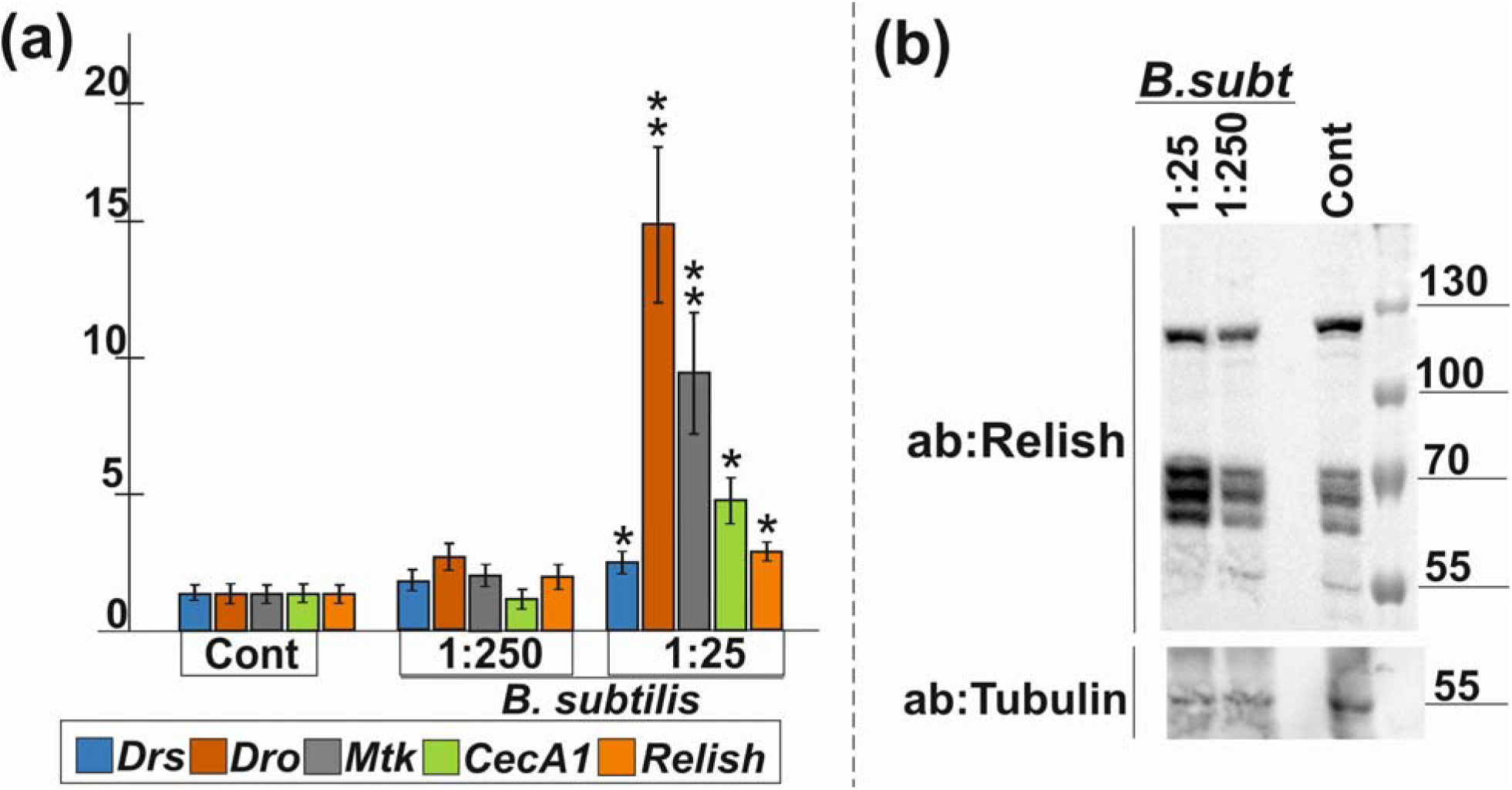
Activation of the immune response to *B. subtilis* in S2 cells. (a) The levels of activation of AMP genes in S2 cells treated with B. subtilis. Cont, no treatment; B. subt, B. subtilis culture diluted 1:250 or 1:25. (b) WB analysis of S2 cell lysates with polyclonal rabbit antibodies against the Relish protein. Three independent experiments were performed. Whiskers show the standard deviation. Asterisks show the statistical significance: (*) P-value ≤ 0.05, (**) P-value ≤ 0.01, or (***) P-value ≤ 0.001 as calculated in the *t*-test.

**Supplementary Table 2.** Relish knockdown has no effect (-) or an effect (+) on gene activation after pathogens in S2 cells.

| Genes | *E. coli* | *M. luteus* | *B. subtilis* | *M. anisopliae* |
| --- | --- | --- | --- | --- |
| *Drs* | + | + | - | - |
| *Dro* | + | + | + | + |
| *Mtk* | + | + | + | + |
| *CecA1* | + | + | + | - |
| *PGRP-SD* | + | + |  |  |
| *PGRP-LB* | - | + |  |  |

## Supplementary List of primers

Primers used in mRNA analysis:

*rp49* transcript,

5΄-ATGACCATCCGCCCAGCATAC and 5΄-GCTTAGCATATCGATCCGACTGG;

*Drosomycin* transcript,

5΄-GTTCGCCCTCTTCGCTGTC and 5΄-GCACACACGACGACAGGTCTC;

*CecropinA1* transcript,

5΄-GCGTTGGTCAGCACACTCG and 5΄-ATCATCGTGGTCAACCTCGG;

*Drosocin* transcript,

5΄-CATCGAGGATCACCTGACTCA and 5΄-TGATCGGATGACTTCTCCG;

*Metchnikowin* transcript,

5΄-GCTACATCAGTGCTGGCAGAG and 5΄-GGTTGGTTAGGATTGAAGGGC;

*PGRP-SA* transcript,

5΄- GGCTGGTGCTACTACTGCTGG and 5΄-CGTGTGATGGATGACCACATAGC;

*PGRP-SD* transcript,

5΄- GATCGGTTTGCTCATCGTTG and 5΄-CCATGCTGTCTATGGCTCCG;

*PGRP-SB1* transcript,

5΄- GATTGAACCACGCAGCAGTTG and 5΄-GGATGTGGAGCAGCCATTAGG;

*PGRP-LE transcript,*

5΄- GCAACTCCACGAACGTCCA and 5΄-CTCGTCCTGATAGTCGTCTTCG;

*PGRP-LB transcript,*

5΄- CCGATCCGACTGGGGTG and 5΄-CGGAGTGGAGTAGCACACGG;

*PGRP-LC transcript,*

5΄- GCAGTACCACCACCCAATCC and 5΄-GGCGATGCTGCCAATACC;

*Relish transcript,*

5΄-CCCACAAACAGCTACAGGAAC and 5΄-TAAAGGCCTCAAAGCAGAGC;

*Dorsal transcript,*

5΄-GTAGTTTCGGAGCCCATCTTCG and 5΄-GAGCAGGATGATCTGGGTGTTG;

*Dif transcript,*

*5΄-GCGCATATGAGCAGCGAACTGACCATC* and

5΄- *GGCGCTCGAGTGGATTCGGGTAGTACTCGA*;

*GNBP1 transcript,*

5΄- CCTGCCAGTCTGCAATCTCG and 5΄- CAGAAGGTTCACAGACGGAGG;

*GNBP3 transcript,*

5΄- GCCTCACCACATCCACCTG and 5΄-GGTGCTCCATTGACCTCGG;

*IMD transcript,*

5΄- CGTGGACGACAACGAACC and 5΄-GCACCACATCCATTGAATTGTAG;

*Spatzle transcript,*

5΄- CAGCCCACGGATGTGAGC and 5΄-CTCGCACTCTTCGATCTGGATG;

*Pelle transcript*

5΄- GGATGGAGCTGCTAGAGAAGCAC and 5΄-CGATCCTGCGGATCCAGG;

*Tube transcript*

5΄- CAATCGCACTGGACATCTGG and 5΄-CTTCCGTTCACCTGCACG;

*Cactus transcript*

5΄- CCGAATGTGCCCAATCTGAC and 5΄-CTGCTCCTCCTGATCCTCTTCTT.

Primers used in ChIP

*Drosomycin* promoter;

5΄-CGACTACGCATCGGCTAAAGC and 5΄-CTCACGGAGCTTGGTAAATGATTTC;

*Cecropin-A1* promoter;

5΄-CAGATGTGTGCTTGGAATCAGAT and 5΄-GATATTGCAGTGAGGTCTGAGCG;

*Metchnikowin* promoter;

5΄-CAATCTGCGACTCGTTTGTCTG and 5΄-GCTCCAAGATTAAGTTGCATCTTAGC;

*Drosocin* promoter;

5΄-CAACCACAGCCAAGACGTG and 5΄-CAGCAGGAAAACGATGGTGAAC;

*PGRP-SA* transcript,

5΄- GGCTGGTGCTACTACTGCTGG and 5΄-CGTGTGATGGATGACCACATAGC;

*PGRP-SD* transcript,

5΄- GATCGGTTTGCTCATCGTTG and 5΄-CCATGCTGTCTATGGCTCCG;

*PGRP-SB1* transcript,

5΄- GATTGAACCACGCAGCAGTTG and 5΄-GGATGTGGAGCAGCCATTAGG;

*PGRP-LE transcript,*

5΄- GCAACTCCACGAACGTCCA and 5΄-CTCGTCCTGATAGTCGTCTTCG;

*PGRP-LB transcript,*

5΄- CCGATCCGACTGGGGTG and 5΄-CGGAGTGGAGTAGCACACGG;

*PGRP-LC transcript,*

5΄- GCAGTACCACCACCCAATCC and 5΄-GGCGATGCTGCCAATACC;

*Relish transcript,*

5΄-CCCACAAACAGCTACAGGAAC and 5΄-TAAAGGCCTCAAAGCAGAGC

*Intergenic spacer;*

5΄-GCCCACAATCGGACATTGAC and 5΄-TCCCACTCCCAAGTCAGGC.

Primers used for the *Relish* dsRNA synthesis

5΄-CGACTCACTATAGGGAGAGACGCTAACGAACGACGATG and

5΄-CGACTCACTATAGGGAGAGACTGATTCAGCAGCGAACAG

